# *De novo* design of a protein fold for small-molecule binding through aromatic π stacking

**DOI:** 10.64898/2026.08.06.743053

**Authors:** Stephanie T. Ouchida, Maggie Horst, Xuxu Gou, Ian Bakanas, A. Katherine Hatstat, Lee Schnaider, Morgan E. Diolaiti, Alan Ashworth, Willliam F. DeGrado

## Abstract

The *de novo* design of proteins that bind chemically complex small molecules has broad chemical and biological implications, but strategies typically rely on a small set of protein scaffolds and require extensive experimental screening. Here, we computationally designed proteins around a minimal aromatic π-stacking motif to bind the anthracycline anticancer drug doxorubicin. Experimental characterization of twelve proteins revealed a µM doxorubicin binder; two additional design cycles improved scaffold stability and binding affinity to yield an 85-residue protein that binds doxorubicin with a dissociation constant of 85 nM. An X-ray crystal structure of the protein-drug complex confirmed the accuracy of the designed π-π stacking interactions. The designed protein could act to protect cultured cells from doxorubicin-induced cytotoxicity. Unlike previous ligand-binding protein designs based on repeat proteins or naturally occurring folds, the designed protein adopts a previously unobserved 5-helix globular fold, indicating that a broader space of folded, functional proteins exists even for compact tertiary structures smaller than 100 residues. These results demonstrate that motif-guided generative protein design can discover compact *de novo* protein folds capable of high-affinity recognition of chemically complex small molecules.

## INTRODUCTION

The *de novo* design of proteins that bind small molecules remains a central challenge in chemistry because successful molecular recognition requires simultaneous specification of both the overall protein scaffold and the atomic-level interactions that define ligand binding.^1–3^ A growing collection of design strategies has enabled the *de novo* generation of proteins that bind increasingly diverse small molecules, with a broad range of structures and functional properties.^4–16^ Many successful binders have been identified through extensive experimental screening,^6,9,11,14,17,18^ whereas only a few studies have reported high computational hit rates.^4,5,7,12^ Despite this progress, experimentally determined structures of *de novo* proteins bound to small molecule ligands remains largely limited to predetermined scaffolds including parameterized helical bundles, repeat proteins^4,7,12,16^ and NTF2-like folds.^5,11^ Extending *de novo* design to discover new protein architectures capable of high-affinity small molecule recognition, therefore, remains an important opportunity for expanding both the scope of molecular recognition and the diversity of functional protein folds.^1,2^

Doxorubicin (1, Fig.1A) is a therapeutically important small molecule whose clinical utility is limited by cumulative, dose-dependent cardiotoxicity.^19–21^ Although liposomal formulations reduce the systemic toxicity of many chemotherapeutics, their relatively large size and heterogeneous biodistribution can limit penetration into target tissues, particularly solid tumors, necessitating the development of alternative drug sequestration and delivery strategies.^22–24^ Compact *de novo* proteins engineered to bind small molecules could provide a complementary platform for these applications owing to their small size, precisely defined molecular composition, and programmable molecular recognition properties.^2,25–27^

**Figure 1.**
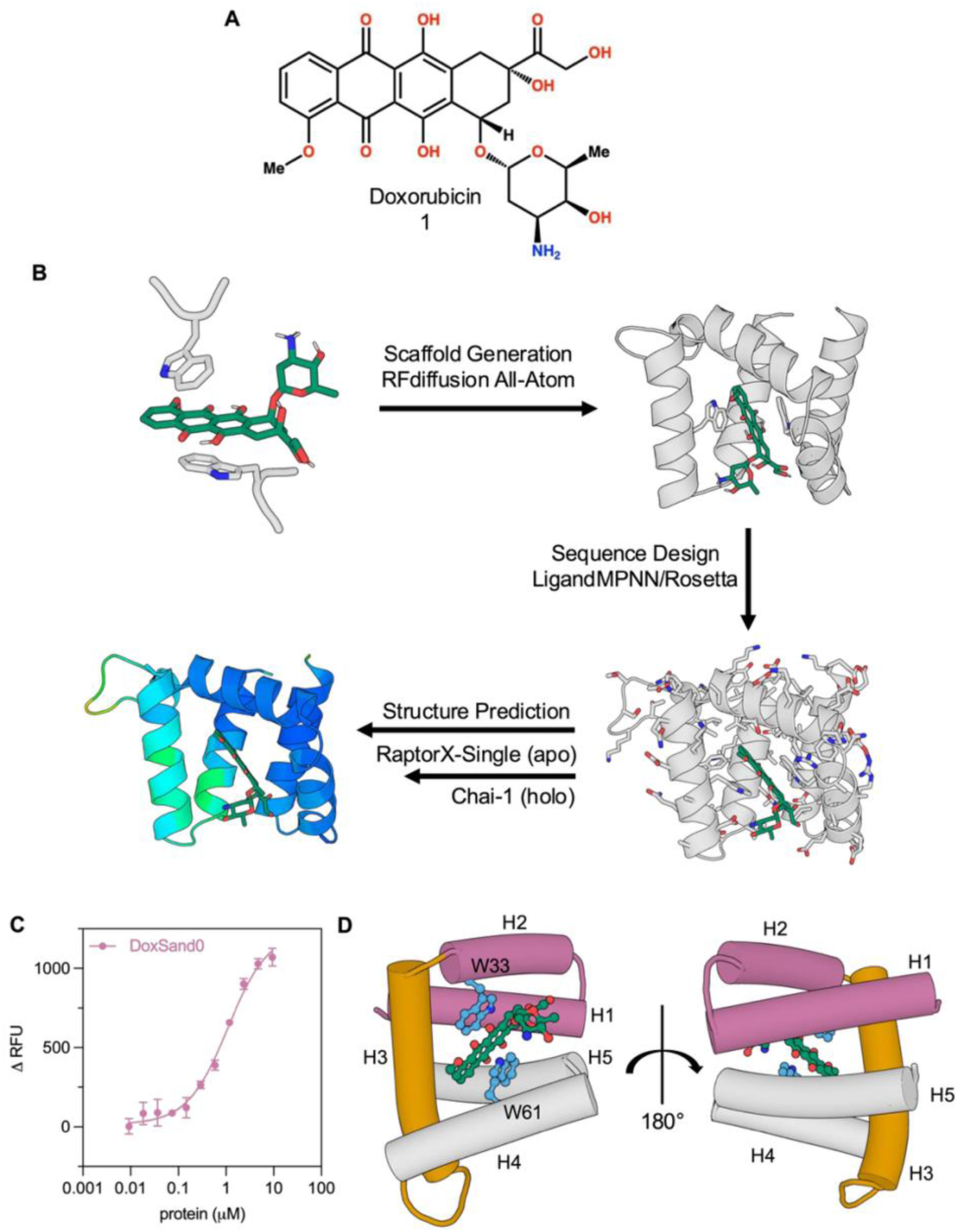
Computational workflow for the design of *de novo* doxorubicin-binding proteins. (A) Chemical structure of doxorubicin. (B) An initial protein–ligand complex containing doxorubicin was used as input for backbone generation with RFdiffusion All-Atom. Protein sequences were designed with LigandMPNN and Rosetta FastRelax, and the resulting designs were evaluated by structure prediction with RaptorX-Single and Chai-1 prior to experimental characterization. (C) Fluorescence titrations of DoxSand0 with doxorubicin. Data were fit to a quadratic binding model, yielding dissociation constants of K_D_= 1.05 ± 0.11 μM for DoxSand0 (best-fit ± parameter SEM from nonlinear regression, n = 3) (D) Chai-1 prediction of DoxSand0. Two views related by a 180° rotation highlight the five-helix scaffold (H1–H5) and the designed doxorubicin-binding pocket.

From a design perspective, doxorubicin is a complex small molecule target. The ligand combines a rigid planar anthracycline scaffold with multiple hydroxyl, carbonyl, and amino substituents, presenting numerous competing opportunities for hydrogen bonding and aromatic stacking without an obvious dominant recognition motif.^28,29^ To facilitate design we prioritized recognition of the anthracycline ring because analysis of crystal structures of anthracycline-bound proteins revealed a recurring interaction in which aromatic residues pack against the planar anthracycline core while surrounding residues accommodate the remaining substituents.^30,31^ We, therefore, expected that pi-pi interactions could establish the overall binding geometry and secondary interactions responsible for affinity and specificity could be optimized independently. Recent advances in generative protein design make this strategy feasible by enabling diverse protein scaffolds to be generated around predefined protein–ligand interaction motifs.^9,13,32^

Here we combine motif-guided binding-site design with RFdiffusion All-Atom,^9^ LigandMPNN,^17^ Rosetta, and structure-prediction-guided filtering to generate *de novo* proteins that bind doxorubicin (Fig. 1B). Starting from an interaction motif derived from a crystal structure of an anthracycline-bound protein, we identified a low-micromolar binder, which could be rapidly optimized to yield an 85-residue protein that binds doxorubicin with nanomolar affinity. These results are noteworthy because there are few examples of the design of small molecule binders which have been validated through the determination of high-resolution structures.^4,5,7,11,12^ Moreover, our results address the question of the extent to which the space of folded, functional structures have been exhaustively sampled during evolution.^33–35^ There are only a limited number of ways that secondary structures can be arranged to create a folded structure, which becomes successively smaller as the size of the protein decreases.^36,37^ While novel structures can be designed in small proteins^38,39^, and small proteins can be designed to bind to protein surfaces,^40–42^ it has been unclear whether nature has sampled the universe of small proteins (< 90 residues) that are stable, cooperatively folded and capable of binding small molecules within a deeply-buried pocket. The design of doxorubicin binders, termed DoxSands, shows that the universe of small, entirely novel miniaturized small molecule proteins has not been exhaustively discovered. This observation is particularly important because proteins of this size can be readily accessed by chemical synthesis, enabling incorporation of noncanonical amino acids and backbone modifications that are difficult to achieve biosynthetically.^43–48^ These capabilities provide opportunities to engineer properties including proteolytic stability, pharmacokinetics, and immune recognition while introducing molecular functions beyond those available to the genetically encoded amino acids.^27,49–54^ Thus, it should be also be possible to achieve entirely new functions that extend well beyond biology.

## RESULTS

### Motif-guided computational protein design identifies a novel fold for binding doxorubicin

To test whether aromatic π-stacking interactions are sufficient to mediate binding of doxorubicin, we generated *de novo* protein scaffolds around a six-residue motif derived from the crystal structure of a protein with moderate affinity for daunomycin, an anthracycline analogue of doxorubicin^30^ (K_D_ = 250 nM, Fig. S1). The motif consists of two three-residue fragments centered on tryptophan residues that position the anthracycline core between opposing aromatic side chains (Fig. 1B). Using this Trp sandwich motif, RFdiffusion All-Atom^9^ was used to generate 500 candidate protein backbones. Scaffolds were initially filtered based on ligand burial and Define Secondary Structure of Proteins (DSSP)^55^-derived assignments, reducing the scaffold pool to 66 candidates. Twelve scaffolds were then selected based on overall structural quality, preservation of the aromatic interaction motif, and accessibility of the designed binding pocket prior to sequence design with LigandMPNN and Rosetta FastRelax^17^ (Fig. S2). Candidate sequences were evaluated by sequential structure prediction. Apo structures were first screened using RaptorX-Single^56^ to eliminate poorly folded designs before ligand-bound complexes were evaluated with Chai-1^57^. Designs were prioritized based on predicted structural confidence, self-consistency with the design model predictions, and preservation of the intended protein–ligand geometry, resulting in a final set of twelve designs for experimental characterization that ranged in length from 93 to 101 residues (Fig. S3).

Nine of the twelve designs showed soluble expression in *E. coli* and were purified for biophysical characterization. One construct, DoxSand0, exhibited measurable binding to doxorubicin with a dissociation constant of K_D_= 1.05 ± 0.11 μM (Fig. 1C). The fold of DoxSand0 appears to be well optimized to serve as a miniaturized protein binder with a jaw-like architecture (Fig. 1D). Starting from the N-terminus, the upper piece of the jaw (colored in pink) is composed of a helical hairpin (H1 and H2), an intervening helix (H3) that forms a hinge (colored in orange), and a C-terminal helical hairpin (H4 and H5) that forms the lower jaw (colored in grey). The “back” two helices (H1 and H5) are tightly interacting, forming the bottom of the cavity, and likely also imparting thermodynamic and structural stability (Fig. 1D, right). The hinge helix (H3) pulls the two front helices (H2 and H4) apart, creating the open cavity that houses the Trp sandwich.

DoxSand0 demonstrated that a motif-guided design strategy could generate a functional doxorubicin binder directly from a small computational design set without experimental affinity maturation. However, biophysical characterization revealed that the scaffold remained incompletely optimized (Fig. 2A-C), prompting structure-guided improvement of the protein’s stability and binding affinity while preserving the designed recognition mechanism.

**Figure 2.**
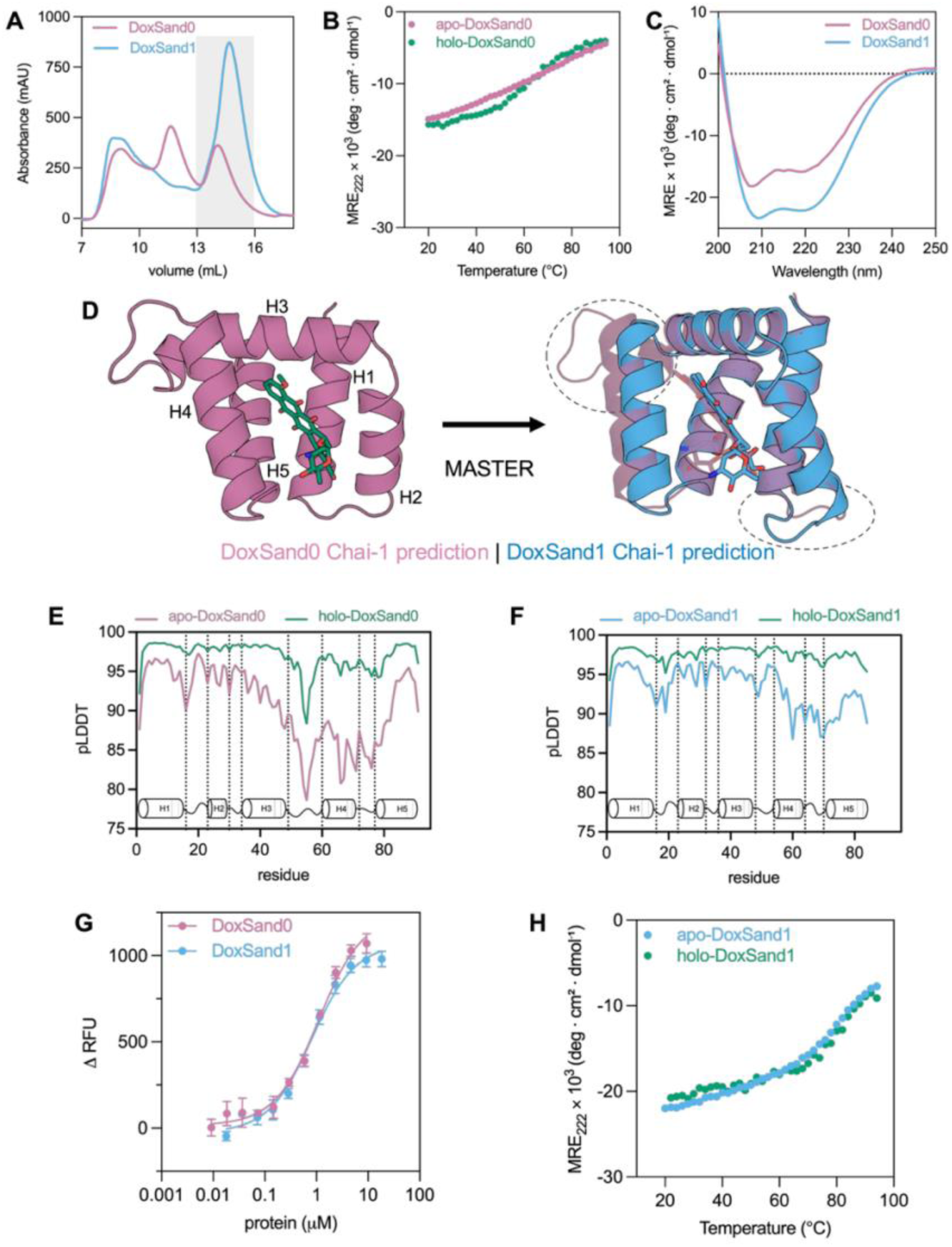
*In silico* scaffold optimization and biophysical characterization of doxorubicin-binding proteins. (A) Size-exclusion chromatography traces of DoxSand0 and DoxSand1. The monomeric fraction used for quantification is indicated by the gray shaded region. (B) Thermal denaturation of apo- and doxorubicin-bound DoxSand0 monitored by circular dichroism at 222 nm. (C) Far-UV circular dichroism spectra of DoxSand0 and DoxSand1. (D) Loop remodeling of DoxSand0 using MASTER generated the DoxSand1 scaffold. Structures shown are Chai-1 predictions; remodeled loop regions are indicated by dashed circles. (E,F) Chai-1-predicted local distance difference test (pLDDT) scores for the apo and holo forms of DoxSand0 (E) and DoxSand1 (F). Secondary structure elements are shown below each plot. (G) Fluorescence titrations of DoxSand0 and DoxSand1 with doxorubicin. Data were fit to a quadratic binding model, yielding dissociation constants of K_D_= 1.05 ± 0.11 μM for DoxSand0 and K_D_= 0.776 ± 0.080 μM for DoxSand1 (best-fit ± parameter SEM, n = 3). (H) Thermal denaturation of apo- and doxorubicin-bound DoxSand1 monitored by circular dichroism at 222 nm.

### Structure-guided scaffold optimization improves stability while preserving ligand recognition

DoxSand0 demonstrated that the motif-guided design strategy successfully generated a functional doxorubicin binder, but biophysical characterization indicated that the scaffold had several limitations. Size-exclusion chromatography revealed that monomeric species constituted only 29% of eluted protein, with the remainder forming higher-order oligomers (Fig. 2A). Consistent with this observation, circular dichroism spectroscopy showed limited cooperative thermal unfolding, suggesting that the designed fold lacked the stability required for a well-defined monomeric scaffold (Fig. 2B,C).

Because the binding site was already functional, we sought to improve the scaffold while preserving the designed ligand-recognition geometry. We hypothesized that the structural heterogeneity arose primarily from backbone geometry. To test this possibility, the loops connecting helices H1–H2 and H3–H4 were remodeled using Method of Accelerated Search for Tertiary Ensemble Representatives (MASTER)^58^ to identify structurally compatible fragments from experimentally determined protein structures while maintaining the surrounding ligand-binding architecture (Fig. 2D). The redesigned scaffold contained 85 amino acids, reducing the overall protein size while preserving the intended binding pocket.

The redesigned scaffold was subsequently optimized through three cycles of LigandMPNN sequence design and Rosetta FastRelax refinement. During sequence design, 31% of the residues were held fixed to preserve the optimized scaffold geometry and ligand-binding motif (Fig. S4). The resulting sequences were evaluated using the same RaptorX-Single and Chai-1 filtering procedure employed in the initial design campaign. Structure prediction indicated increased confidence throughout the remodeled loop regions (Fig. 2E,F). Characterization of DoxSand1 by size exclusion chromatography showed that the monomeric fraction increased from 29% to 51% (Fig 2A). DoxSand1 also showed improved thermal stability while maintaining a similar binding affinity to doxorubicin, with K_D_=0.78±0.08 μM (comparable to the starting DoxSand0, which had a K_D_=1.05±0.11; Fig. 2C,G,H). These results indicate that scaffold stability could be substantially improved without disrupting the designed ligand-recognition mechanism, suggesting that scaffold optimization and binding-site optimization could be pursued independently.

### Binding-site optimization yields a nanomolar-affinity doxorubicin binder

Having established a stable scaffold that binds doxorubicin, we next tested the designed binding mode by individually mutating residues predicted to contact the ligand and evaluating the resulting variants by fluorescence binding assays (Fig. 3A). Mutation of Trp33 revealed a strong preference for a bulky aromatic residue at this position. Substitution with phenylalanine largely preserved binding (K_D_= 2.18 ± 0.17 μM), whereas tyrosine (K_D_ > 10 μM) and alanine (K_D_ > 20 μM) substantially reduced affinity (Fig. 3B). Trp61 was more tolerant to substitution, with phenylalanine (K_D_= 1.03 ± 0.10 μM) and tyrosine (K_D_= 7.57 ± 1.45 μM) retaining measurable binding, while alanine (K_D_ > 20 μM) markedly weakened binding (Fig. 3C). These results suggest that favorable packing interactions at both positions are important for doxorubicin binding, with stricter steric requirements at Trp33 than at Trp61 (Fig. 3D). Consistent with the design model, mutational analysis of Trp33, Trp61, and Asp65 supports a binding mode in which the two tryptophan residues form the central ligand-binding interface, while surrounding polar interaction further stabilizes the protein–ligand complex. In contrast, mutation of His73 to alanine unexpectedly increased affinity, yielding a dissociation constant of 95 ± 31 nM (Fig. 3E). The Chai-1 predictions for the H73A variant suggest a subtle but meaningful rearrangement of the binding site, including a slight shift in the ligand pose and a corresponding change in the Trp61 rotamer, which together improve ligand burial and the geometry of the aromatic interaction network (Fig. 3F).

**Figure 3.**
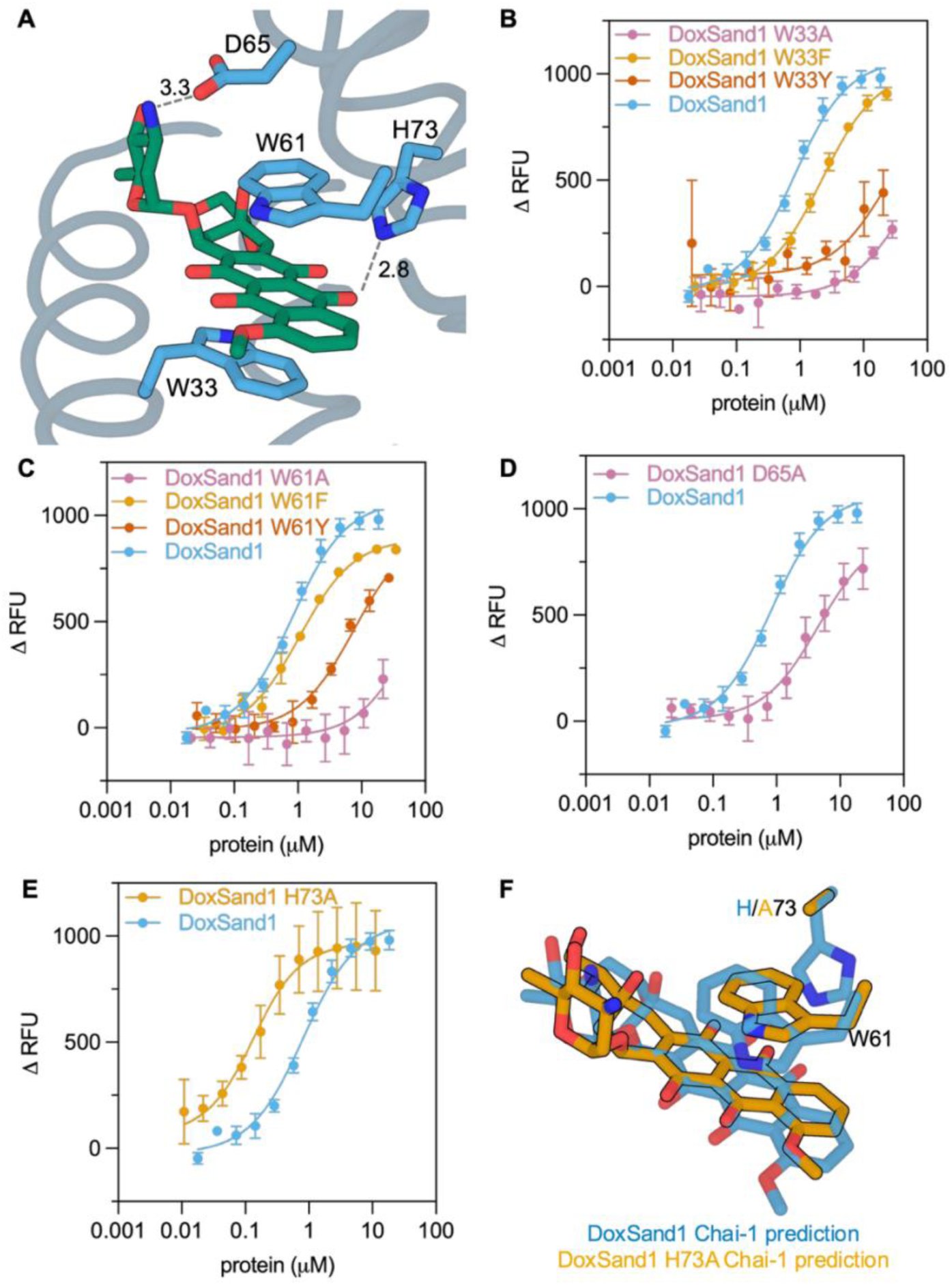
Mutational analysis of the DoxSand1 binding pocket. (A) Predicted binding-site interactions between DoxSand1 and doxorubicin. Residues selected for mutational analysis are shown as sticks. Hydrogen-bonding interactions are shown as dashed lines, with heavy atom–heavy atom (donor–acceptor) distances indicated in angstroms. (B–E) Fluorescence titrations of DoxSand1 variants with doxorubicin. Data were fit to a quadratic binding model. Dissociation constants (K_D_) were: W33A, > 20 μM; W33Y, > 10 μM; W33F, 2.18 ± 0.17 μM (B) W61A, > 20 μM; W61F, 1.03 ± 0.10 μM; W61Y, 7.57 ± 1.45 μM (C); D65A, 4.51 ± 1.06 μM (D); H73A, 95.0 ± 31.0 nM (E). (best-fit ± parameter SEM from nonlinear regression, n=3). Variants W33A, W33Y, and W61A exhibited substantially reduced binding but did not reach saturation over the concentration range tested, preventing reliable determination of K_D_. (F) Overlay of Chai-1 predictions for DoxSand1 (blue) and the H73A mutant (orange), showing the predicted binding-site architecture

Despite its improved binding affinity, the H73A variant exhibited a reduced monomeric fraction (36%) compared to DoxSand1 (51%) by size-exclusion chromatography (Fig. 4A), suggesting that removal of the His73 side chain compromised scaffold stability even as it enhanced ligand recognition. To restore stability while maintaining the improved binding properties of the H73A variant, we introduced a structure-guided V46I substitution adjacent to the binding pocket; the increased bulk of the Ile relative to the initial Val sidechain was selected to improve van der Waals packing by filling a small cavity at the helix-helix interface as well as increasing the hydrophobic driving force for folding. Because H73A emerged as the lead variant, we subsequently recharacterized its affinity using fluorescence titrations spanning three doxorubicin concentrations and globally fit the resulting datasets with a shared dissociation constant, yielding a more robust affinity estimate of K_D_ = 58 ± 17 nM (Fig. 4C). The resulting construct, DoxSand2, exhibited an increased monomeric fraction (61%), retained nanomolar affinity for doxorubicin (K_D_ = 85 ± 16 nM) and displayed improved biophysical properties relative to DoxSand1 H73A (Fig. 4D,E-G). Thus, through iterative structure-guided optimization, we transformed the initial computational hit into a higher-affinity binder with improved biophysical properties while preserving the designed binding mode.

**Figure 4.**
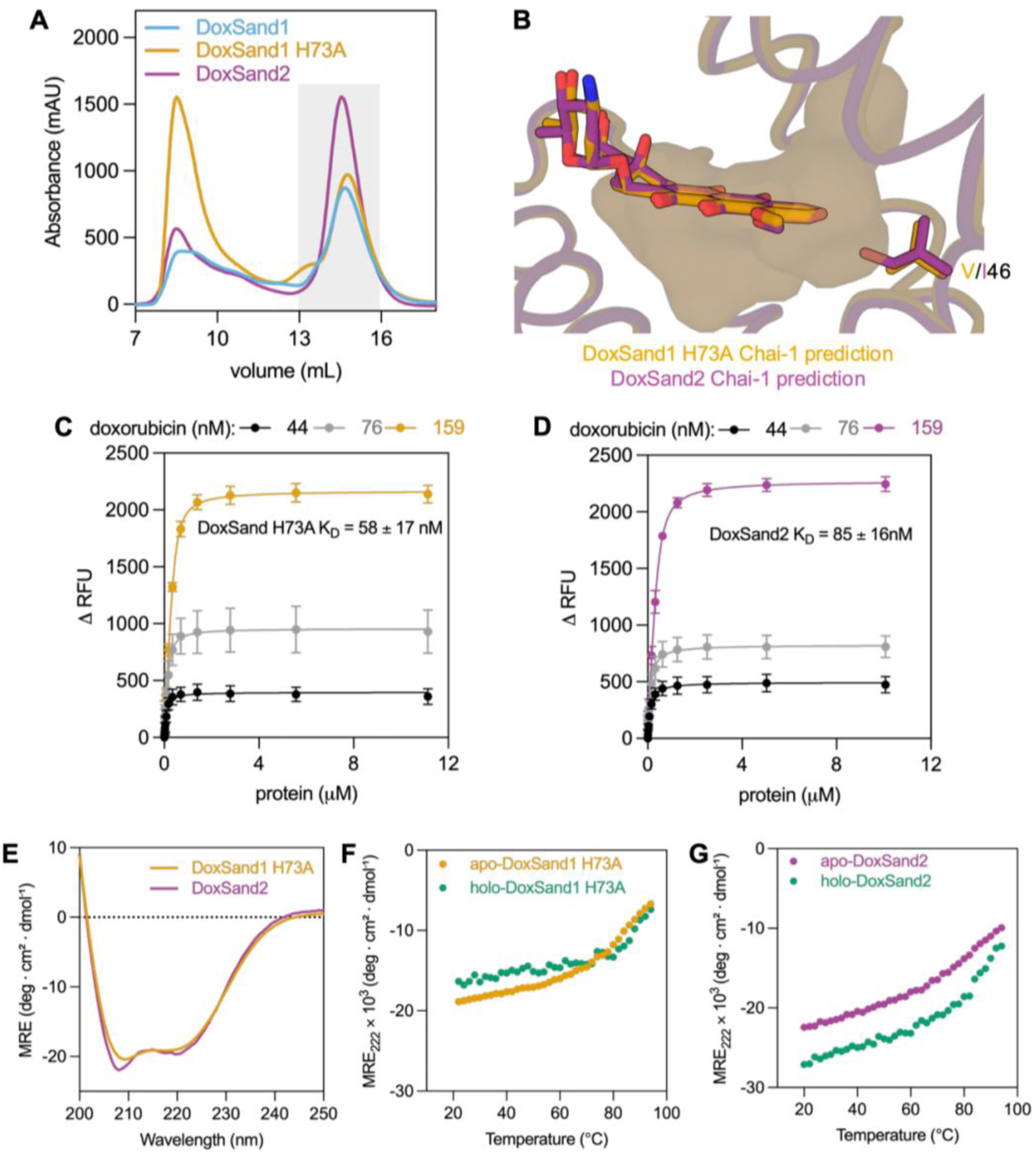
Structure-guided optimization of the DoxSand scaffold. (A) Size-exclusion chromatography traces of DoxSand1, DoxSand1 H73A, and DoxSand2. Gray shaded regions denote the monomeric fractions used for subsequent assays, representing 51% (DoxSand1), 36% (DoxSand1 H73A), and 61% (DoxSand2) of the total eluted protein. (B) Overlay of Chai-1 predictions for DoxSand1 H73A (orange) and DoxSand2 (magenta), showing that the V46I substitution fills an unoccupied space within the doxorubicin-binding pocket, resulting in improved predicted packing around the ligand. (C,D) Fluorescence titrations of DoxSand1 H73A (C) and DoxSand2 (D) with 44, 76, and 159 nM doxorubicin. Data were globally fit by nonlinear regression using a quadratic binding model with a shared dissociation constant, yielding K_D_= 58 ± 17 nM for DoxSand1 H73A and K_D_= 85 ± 16 nM for DoxSand2 (best-fit ± parameter SEM from nonlinear regression, n = 3). (E) Far-UV circular dichroism spectra of DoxSand1 H73A and DoxSand2. (F,G) Thermal denaturation monitored by circular dichroism for apo- and doxorubicin-bound DoxSand1 H73A (F) and DoxSand2 (G).

### Crystal structure validates the designed recognition mechanism

To evaluate the accuracy of the computational design, we determined the crystal structure of the doxorubicin-bound DoxSand2 complex at 1.56 Å resolution (Fig. 5). The experimentally determined structure agrees closely with the design model, with an overall Cα RMSD of 0.57 Å relative to the Chai-1 prediction (Fig. 5A). The crystal structure confirms that DoxSand2 adopts the desired fold: a compact five-helix topology composed of an N-terminal helix-hairpin (H1-H2), a central helix (H3), and a second helix-hairpin (H4-H5) that packs across the central helix to form a tightly integrated tertiary structure (Fig. 5A). The helices position the designed ligand-binding pocket at the interface between the two helix-hairpin motifs.

**Figure 5.**
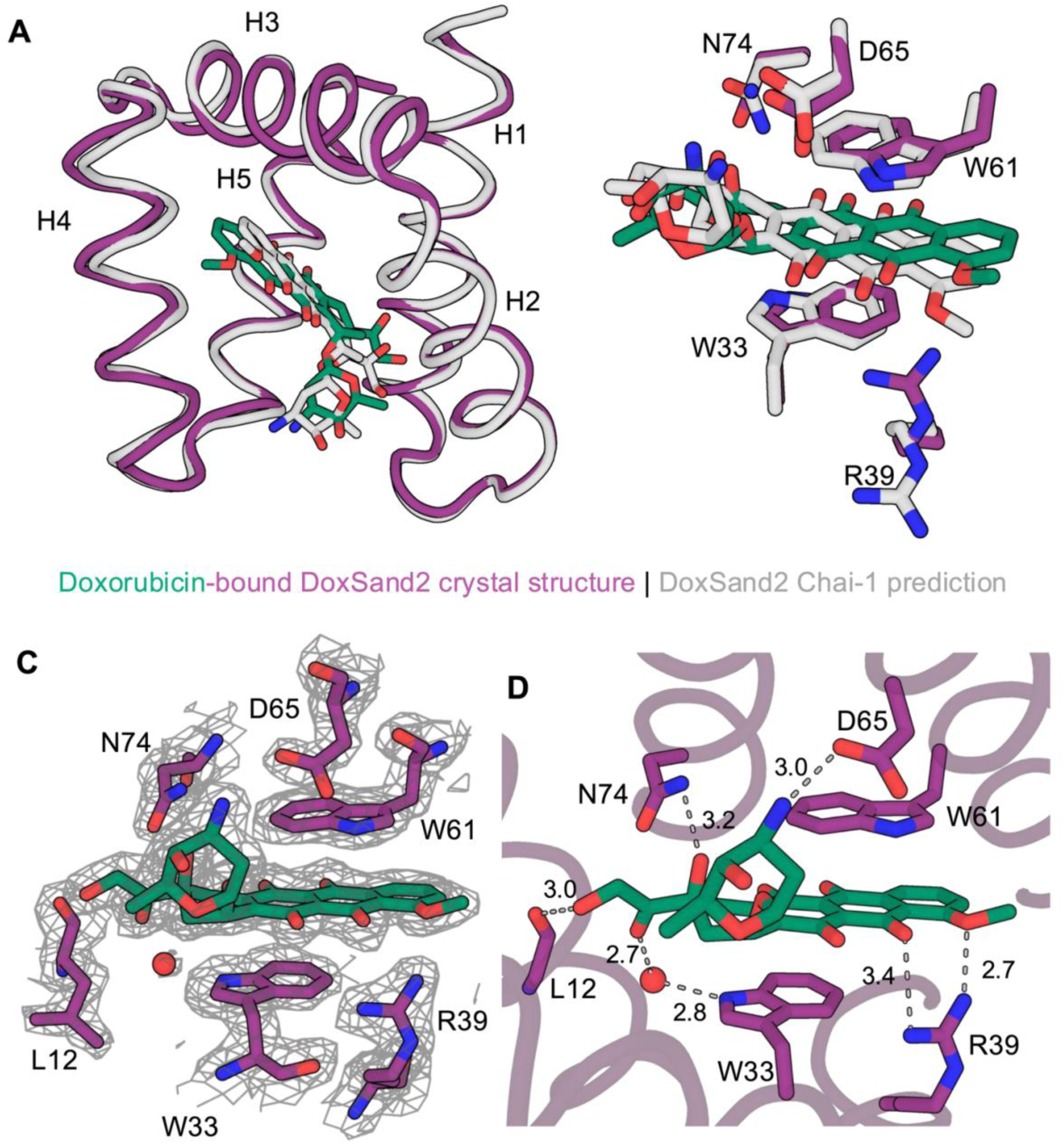
Crystal structure of the doxorubicin-bound DoxSand2 complex. (A) Superposition of doxorubicin-bound DoxSand2 crystal structure (doxorubicin colored in dark green, DoxSand2 colored in magenta) and the DoxSand2 Chai-1 prediction (doxorubicin and DoxSand2 colored gray). Doxorubicin is shown as sticks. (B) Overlay of the binding-site residues from the crystal structure and Chai-1 prediction after backbone superposition. (C) Binding pocket of the doxorubicin-bound DoxSand2 crystal structure. The 2Fo−Fc composites omit electron density maps for doxorubicin and surrounding binding-site residues is contoured at 1σ. (D) Binding-site interactions observed in the crystal structure. Hydrogen-bonding interactions are shown as dashed lines, with heavy atom–heavy atom (donor–acceptor) distances indicated in angstroms. Ordered water molecule is shown as red sphere.

Inspection of the binding pocket further confirmed that the principal design features were preserved. The anthracycline core of doxorubicin is positioned between the two designed tryptophan residues (W33 and W61), reproducing the aromatic sandwich interaction that served as the foundation for the computational design (Fig. 5B). Likewise, Asp65, identified by mutational analysis as an important determinant of binding, occupies the predicted position adjacent to the amino sugar of doxorubicin and participates in the expected interaction. The agreement between the mutational analysis and the crystal structure demonstrates that the dominant interactions responsible for ligand recognition were accurately encoded during computational design.

Comparison of the crystal structure with the Chai-1 model also shows that the ligand is predicted with good accuracy (ligand RMSD = 1.34 Å), with only a small, angstrom-level shift of the ligand in the binding pocket (Fig. 5A,B). As in the model, the anthracycline core is sandwiched between Trp33 and Trp61 in the designed aromatic recognition geometry. A small shift in the ligand position establishes several interactions absent from the design model, including a direct hydrogen bond between the doxorubicin hydroxyl group and the backbone carbonyl of Leu12, as well as a direct interaction with Arg39. In addition, an ordered water molecule bridges the doxorubicin carbonyl group and Trp33, creating a water-mediated interaction that was not explicitly modeled during design which did not consider solvent(Fig. 5C,D).

To assess the structural novelty of DoxSand2, the experimentally determined structure was searched against both the PDB100 and AlphaFold/UniProt50 databases using Foldseek (3Di/AA mode).^59^ No statistically significant structural homologs were identified in either database. Searches against PDB100 yielded only low-confidence matches (9–26% sequence identity; Foldseek probability <0.3; E-value >1) (Fig. S5A). Similarly, searches against AlphaFold/UniProt50 identified no statistically significant homologs; although 24 hits exhibited Foldseek probabilities ≥0.5, all retained E-values >1 (Fig. S5B). Moreover, the highest-scoring matches were distributed across diverse α-helical proteins, with no enrichment for a single structural family. These results indicate that DoxSand2 adopts an α-helical architecture that is distinct from previously characterized protein scaffolds.

### Designed binders attenuate doxorubicin cytotoxicity in COV362 cells

To determine whether the designed proteins modulate the biological activity of doxorubicin, we evaluated their effects on the proliferation of COV362 ovarian cancer cells (Fig. 6). Dose–response curves were first established using free doxorubicin and then repeated using doxorubicin preincubated with either DoxSand1 or DoxSand2, each at a fixed protein concentration of 20 µM. Preincubation with DoxSand1 increased the half-maximal inhibitory concentration (IC₅₀) of doxorubicin approximately 17-fold, from 6.9 to 116 nM. DoxSand2 produced an even larger effect, increasing the apparent IC₅₀ to 951 nM, corresponding to a ∼140-fold shift relative to free doxorubicin. The greater protective effect of DoxSand2 is consistent with its higher affinity for doxorubicin relative to DoxSand1. These results demonstrate that the designed proteins effectively sequester doxorubicin in a cellular environment and substantially attenuate its cytotoxic activity.

**Figure 6.**
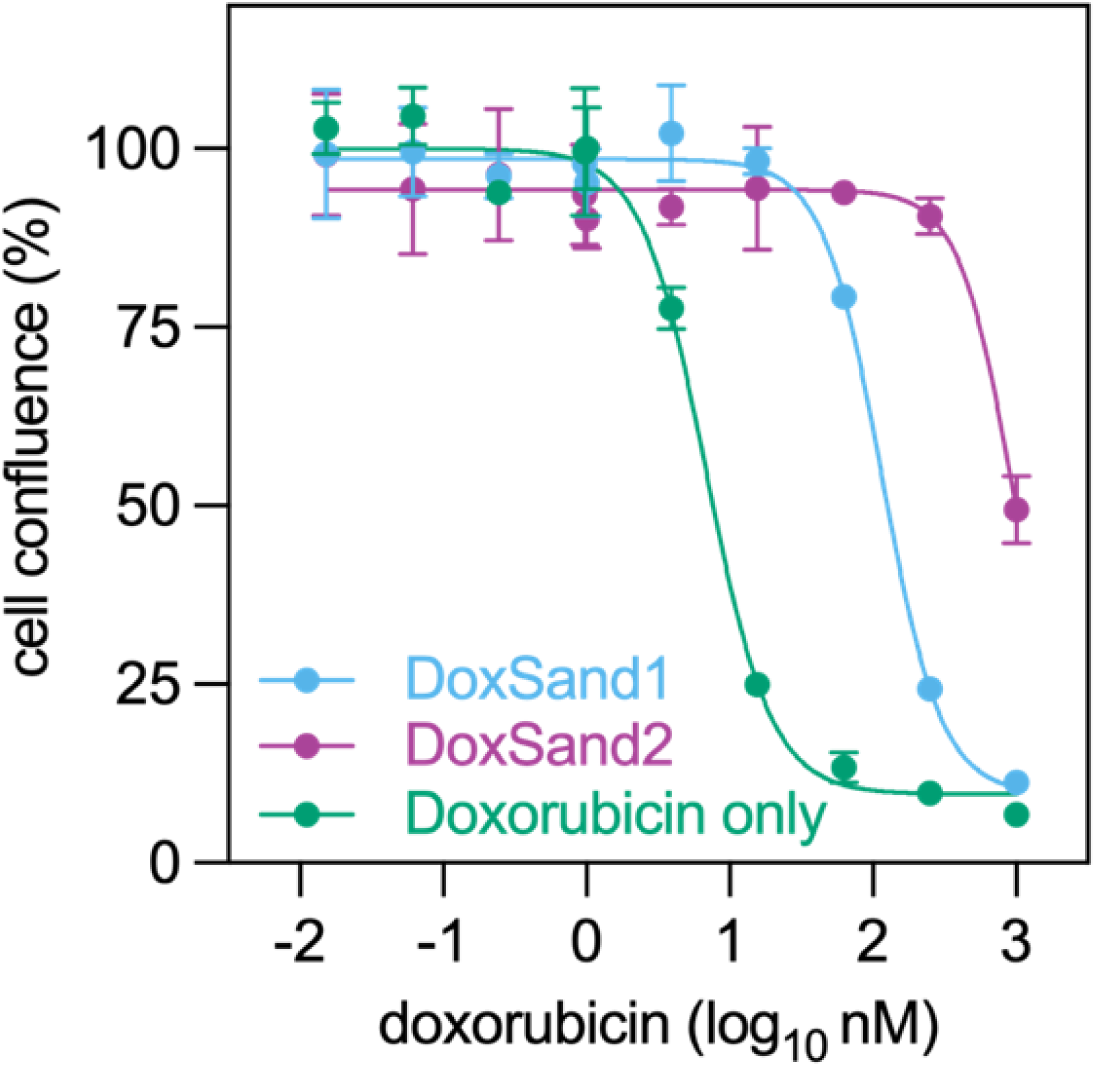
DoxSand proteins reduce doxorubicin cytotoxicity in COV362 cells. COV362 cell confluence following treatment with increasing concentrations of free doxorubicin or doxorubicin preincubated with 20 μM DoxSand1 or DoxSand2. Curves represent fits to a four-parameter dose–response model. Data are shown as mean ± SD (n = 2). The apparent IC₅₀ values were 6.85 nM (95% CI, 5.80–8.38 nM) for free doxorubicin, 115.7 nM (95% CI, 95.6–139.6 nM) for DoxSand1, and 950.7 nM (95% CI, 805.8–1122 nM) for DoxSand2.

## DISCUSSION

We designed a compact protein architecture *de novo* that binds a chemically complex therapeutic molecule with nanomolar affinity. Though only 85 residues long, the protein adopts a well-defined five-helix topology that accurately positions doxorubicin within a buried ligand-binding pocket. Most structurally characterized *de novo* proteins bound to small molecule ligands have adopted a relatively limited set of scaffold architectures, primarily parameterized helical bundles and proteins based on the NTF2 fold. (Table S1).^4,5,7,11^ In contrast, DoxSand2 adopts a distinct five-helix topology organized around two helical hairpins connected by a central hinge helix which produces a jaw-like binding cavity that encloses the ligand while maintaining an exceptionally compact scaffold. This organization demonstrates that compact ligand-binding cavities can be supported by substantially different tertiary architectures than those represented among current *de novo* binders. More broadly, these findings suggest that the current collection of structurally characterized *de novo* small-molecule binders likely underrepresents the diversity of protein architectures capable of supporting high-affinity molecular recognition, even among proteins smaller than 90 residues.

A key aspect of this work is the use of an aromatic π-stacking interaction to guide binder design. Unlike many previous *de novo* binder design efforts which optimize extensive interaction networks within predefined scaffold architectures,^4,5,7,11,12^ we asked whether an aromatic π-stacking sandwich interaction could seed the discovery of an entirely new ligand-binding fold. The resulting nanomolar binder demonstrates that establishing a dominant geometric interaction can substantially reduce the complexity of designing binders for chemically complex small molecules while leaving secondary interactions to be optimized during subsequent refinement.

The crystal structure provides an opportunity to evaluate the current capabilities of ligand-aware computational design.^9,17,57^ The experimentally determined protein closely matches the design model, whereas the bound ligand adopts a shifted pose that preserves the designed π-stacking interaction while introducing additional hydrogen-bonding interactions and ordered water molecules not explicitly modeled during design. These observations indicate that current computational methods accurately specify overall protein architecture and the dominant geometric features governing molecular recognition while highlighting opportunities to improve prediction of ligand placement and explicit treatment of solvents. More broadly, the progression from an initial computational hit to a nanomolar binder demonstrates that computational design can provide experimentally useful starting points for rational optimization without requiring extensive empirical screening.^4,5,7,12^ The ability of DoxSand2 to reduce doxorubicin cytotoxicity demonstrates that compact *de novo* binders can meaningfully modulate the activity of clinically relevant small molecules, illustrating the functional potential of these designed proteins beyond structural validation.^7,27^

## CONCLUSIONS

A minimal aromatic π-stacking motif was sufficient to seed the *de novo* design of a previously unobserved protein fold that binds a chemically complex small molecule with nanomolar affinity. These findings demonstrate that compact ligand-binding proteins need not be limited to naturally occurring folds or previously established design architectures. DoxSand demonstrates that the structural diversity of functional ligand-binding proteins remains far from fully explored.

## ACKNOWLEDGEMENTS

The authors thank Andrew Ecker for insightful discussions throughout this work. The structure of DoxSand2 was determined using diffraction data collected at synchrotron X-ray beamline 8.3.1 of the Advanced Light Source, a DOE Office of Science User Facility under Contract No. DE-AC02-05CH11231, supported in part by the ALS-ENABLE program funded by the National Institutes of Health, National Institute of General Medical Sciences, grant P30 GM124169-01. The authors thank Mehagan Hopkins and James Holton for critical beamline support. M.H. and I.B. were supported by a Ruth L. Kirschstein NRSA Postdoctoral Fellowship from the National Institute of General Medical Sciences of the National Institutes of Health under award number F32GM163375 and F32GM154484, respectively. L.S. and A.K.H. were supported by National Institute of General Medical Sciences of the National Institutes of Health under award number F32GM143869-01A1 and K99GM155611, respectively. X.G. was supported by The Baker Street Foundation. Research reported in this publication was primarily supported by the National Institute of General Medical Sciences of the National Institutes of Health under award number R35GM-122603 and by the National Science Foundation under award number MCB-2306190. The content is solely the responsibility of the authors and does not necessarily represent the official views of the National Institutes of Health. The authors used ChatGPT (OpenAI) to assist with improving the clarity and grammar of portions of the manuscript. The authors reviewed, edited, and take full responsibility for the final content.

