## Supplemental Methods and Information for "*De novo* design of a protein fold for small-molecule binding through aromatic π stacking"

Affiliations:
[1] Department of Pharmaceutical Chemistry & Cardiovascular Research Institute, University of California, San Francisco, CA 94158, USA.
[2] Department of Molecular and Chemical Life Sciences, Tohoku University, Sendai, Japan
[3]Helen Diller Family Comprehensive Cancer Center, University of California at San Francisco, San Francisco, CA 94158, USA

### SUPPLEMENTAL METHODS

*Overview of Computational Design Process*

An aromatic sandwich motif was extracted from the crystal structure of a daunorubicin-binding protein complex (PDB ID: 3F8F). The motif consisted of two tryptophan-containing three-residue fragments that form π-stacking interactions with the anthracycline core of daunorubicin. Doxorubicin was positioned by pair-fitting its anthracycline core onto the daunorubicin coordinates from the crystal structure, and the resulting protein–ligand motif was used as a fixed input for scaffold generation.

Protein backbones were generated using RFdiffusion All-Atom[1] with scaffold lengths ranging from 100 to 120 residues. Scaffolds were filtered based on ligand burial and secondary structure content. The remaining scaffolds were subjected to sequence design using LigandMPNN followed by Rosetta FastRelax (RosettaCommons, Linux binary release 315) refinement.[2] Structures were evaluated using RaptorX-Single[3] and subsequently analyzed using Chai-1 (version 0.5.2)[4] holo structure prediction. Designs were manually selected for experimental characterization based on preservation of the aromatic sandwich geometry and consistency of ligand placement across multiple Chai-1 predictions.

Following identification of the initial binder DoxSand0, a second round of design was performed to improve scaffold stability and monomeric stability through loop replacement using MASTER[5]. Redesigned scaffolds were subsequently subjected to the same sequence design and structure prediction workflow.

Custom Python scripts were developed to implement the computational protein design workflow and perform downstream structural analyses. ChatGPT (OpenAI) was used as an interactive programming assistant to assist with code generation, debugging, and refactoring during pipeline implementation and data analysis. All computational workflows, parameter selection, analyses, and interpretation of results were designed, validated, and verified by the authors.

Representative inputs from design pipeline are shown below:

RFdiffusion all-atom inference command:

```
run_inference.py \
     inference.deterministic=True \
     diffuser.T=100 \
     inference.output_prefix=<output_prefix> \
     inference.input_pdb= <input_pdb> \
     contigmap.contigs=['100-150'] \
     inference.ligand=DM2 \
     inference.num_designs=1 \
     inference.design_startnum=0
```

LigandMPNN run command:

```
LigandMPNN/run.py \
     --model_type "ligand_mpnn" \
     --seed 111 \
     --pdb_path "input_pdb_path" \
     --out_folder "output_path" \
     --batch_size 2 \
     --number_of_batches 5 \
     --fixed_residues "lmpnn_fixed_resis" \
     --omit_AA "CM" \
```

Rosetta FastRelax run command:

```
rosetta.binary.linux.release-
315/main/source/bin/relax.static.linuxgccrelease \
    -database {rosetta_path}/main/database/ \
     -in:file:s $input_pdb \
     -extra_res_fa {ligand_params_file} \
     -relax:quick \
     -constraints:cst_fa_file {constraints_file} \
        -constraints:cst_fa_weight 5.0 \
     -out:no_nstruct_label \
```

RaptorX-Single run command:

```
RaptorX-Single/pred.py \
     fasta_path \
     params/RaptorX-Single-ESM1b-ESM1v-ProtTrans.pt \
     --out_dir={output_directory_path} \
     --device_id=0
```

Chai1 run

```
chai1_lab.chai1.run_inference(
     fasta_file=fasta_path,
     output_dir=output_dir,
```

```
num_trunk_recycles=3,
num_diffn_timesteps=200,
seed=42,
device=torch.device("cuda:0"),
use_esm_embeddings=True)
```

*Scaffold Generation*

Protein backbones were generated using RFdiffusion All-Atom with scaffold lengths ranging from 100 to 120 residues. The input motif was derived from the daunorubicin-binding protein structure (PDB ID: 3F8F) and consisted of two fixed three-residue fragments containing the tryptophan residues responsible for aromatic stacking interactions with the anthracycline core. Doxorubicin coordinates were generated by pair-fitting the anthracycline core of doxorubicin onto the daunorubicin molecule present in the crystal structure.

A total of 500 scaffolds were generated. Scaffolds were filtered based on ligand burial and secondary structure content. Ligand burial was quantified using solvent-accessible surface area (SASA) calculations performed with the Shrake–Rupley algorithm implemented in Biopython using a probe radius of 1.0 Å. Scaffolds were required to exhibit a ligand ΔSASA greater than 220 $Å^2$ upon binding. Secondary structure assignments were determined using DSSP through Biopython, and scaffolds containing more than six consecutive residues assigned as loop were removed. Application of these filters reduced the design pool from 500 to 66 scaffolds. The 66 filtered scaffolds were manually evaluated in PyMOL for overall scaffold geometry, preservation of the aromatic interaction motif, and accessibility of the ligand-binding pocket. Based on this assessment, 12 scaffolds were selected for subsequent iterative sequence design.

*Sequence Design and Structural Filtering*

The 12 selected scaffolds were subjected to three iterative rounds of fixed-backbone sequence design using LigandMPNN, each followed by backbone relaxation with Rosetta FastRelax. During sequence design, the aromatic sandwich residues (W33 and W61) were held fixed. A total of 512 sequences were generated for each scaffold, affording 6144 sequences. Designed sequences were refined using Rosetta FastRelax with ligand movement enabled during relaxation. Apo structures were subsequently predicted using RaptorX-Single and filtered based on average predicted confidence (pLDDT > 85), backbone RMSD relative to the design model (<1.5 Å), and net charge between −3 and −7. Application of these criteria reduced the design pool to 170 sequences.

*Chai-1 Structure Prediction and Design Selection*

The remaining sequences were evaluated using Chai-1 holo structure prediction. Five independent Chai-1 predictions were generated for each sequence using doxorubicin represented by the SMILES string

```
C[C@H]1[C@H]([C@H](C[C@@H](O1)O[C@H]2C[C@@](Cc3c2c(c4c(c3O)C(=O)
c5cccc(c5C4=O)OC)O)(C(=O)CO)O)N)O
```

Predicted structures were analyzed using custom scripts and manually inspected. Designs were prioritized based on preservation of the aromatic sandwich geometry, consistency of ligand placement, and agreement among the five Chai-1 predictions. Emphasis was placed on maintaining the orientation of the tryptophan sandwich residues and the self-consistency of the predicted ligand pose across all five predictions. Chai-1 predictions were used for ranking and design selection; no additional computational filtering was applied at this stage. Twelve sequences derived from six scaffolds were selected for experimental characterization.

For the final DoxSand2 structure prediction used for comparison with the experimentally determined crystal structure, Chai-1 was run using an alternative doxorubicin SMILES representation,

```
COc1cccc2C(=O)c3c(O)c4C[C@](O)(C[C@H](O[C@H]5C[C@H](N)[C@H](O)[C
@H](C)O5)c4c(O)c3C(=O)c12)C(=O)CO
```

because this representation consistently yielded Chai-1 predictions with the correct stereochemistry of doxorubicin.

*Loop Remodeling*

Following experimental characterization of the initial binder DoxSand0, a second round of design was performed to improve scaffold stability and monomeric stability. Loop replacement was performed using MASTER. Two six-residue helical fragments flanking each loop were used as search queries against structures in the Protein Data Bank. Loop lengths ranging from 3 to 10 residues were sampled using RMSD cutoffs of 1.0–1.5 Å. The wgap option in MASTER was used to identify loops of the specified length connecting the two helices within the defined RMSD threshold. The resulting loop candidates were clustered based on RMSD and ranked according to cluster size. Two loop regions, connecting helices H1–H2 and H3–H4, were replaced with structurally compatible fragments identified through this MASTER search procedure. The redesigned scaffold was reduced from 92 residues to 85 residues while preserving the binding-site architecture. Redesigned structures were subjected to three iterative rounds of fixed-backbone sequence design with LigandMPNN, during which 31% of scaffold residues were held fixed, each followed by Rosetta FastRelax refinement, with Rosetta constraints applied to residues W33 and W61 relative to the bound doxorubicin to preserve the binding-site geometry during sequence optimization. Refined designs were subsequently filtered using RaptorX-Single and evaluated with Chai-1 as described above.

*Structural similarity search*

Structural similarity searches were performed using the Foldseek[6] web server in 3Di-AA search mode with default settings. The holo monomer structure of DoxSand2 (PDB: 36IQ) was used as the query model. Searches were performed against the PDB100 and AFDB50 databases using default parameters. Hits were evaluated based on Foldseek probability, E-value, and sequence identity.

*Protein expression and purification*

Gene fragments encoding the designed proteins followed by TEV protease cut site (sequence: ENLYFQS) and C-terminal 6×His tag were obtained from Twist Bioscience and cloned into pET28a(+) by Gibson assembly. Plasmid constructs were verified by Sanger sequencing (Azenta Life Sciences) using the T7 primer. Plasmids were transformed into *E. coli* BL21(DE3) cells and grown in Luria Broth (Miller formulation) medium supplemented with kanamycin to final concentration of 50 µg/mL at 220 rpm, 37 °C. Cultures were induced with Isopropyl β-D-1-thiogalactopyranoside (IPTG) to a final concentration of 1 mM at an $OD_{600}$ of 0.6–0.8 and grown for an additional 4 hours at 220 rpm, 37 °C before harvesting by centrifugation.

Cell pellets were resuspended in 1× phosphate buffered saline (PBS), pH 7.4 supplemented with 20 mM imidazole and lysed by sonication (Sonic Dismembrator Model 500, Fisher Scientific) before clarification by centrifugation. Clarified lysates were loaded onto Ni-NTA resin (HisPur™ Ni-NTA Resin, ThermoFisher) equilibrated 1x PBS, 20mM imidazole and washed with 1× PBS, 20 mM imidazole. Bound proteins were eluted with 1× PBS containing 250 mM imidazole. Eluted fractions were concentrated and further purified by size exclusion chromatography on a Superdex 75 Increase 10/300 GL column equilibrated in 1× PBS (pH 7.4). Protein purity was assessed by SDS-PAGE (Invitrogen NuPAGE 4-12% Bis-Tris Mini Protein Gel, ThermoFisher) and concentrations were determined by UV–visible spectroscopy. Untagged (e.g. hexahistidine tag removed by TEV cleavage) and $His_6$-tagged proteins showed no significant difference in binding; therefore, His-tagged protein was used for experiments.

*Determination of binding dissociation constants*

Fluorescence quenching experiments were performed in black, non-binding surface (NBS) 384-well microplates (Corning®, Cat. No. CLS3575BC) with 1X PBS, pH 7.4 as the assay buffer. Protein solutions were prepared at an initial target concentration of ~10 µM and serially diluted in a 96-well plate (100 µL per well). Exact protein concentrations were determined by UV–visible spectroscopy and used for all subsequent calculations. Aliquots (85 µL) of each protein dilution were transferred to the 384-well plate. Doxorubicin hydrochloride (Cayman Chemical, Cat. No. 15007) was dissolved in dimethyl sulfoxide (DMSO) to prepare a 20 mM stock solution and stored at –20 °C. For fluorescence quenching assays, a working stock solution was prepared by diluting the DMSO stock into Milli-Q water. The concentration of the working stock was verified by UV–visible spectroscopy using an extinction coefficient of 13,500 $M^{-1}$ $cm^{-1}$ at 480 nm.[7] Unless otherwise indicated, the working stock was prepared to yield a final doxorubicin concentration of 100 nM upon addition of 5 µL to each well (final assay volume, 90 µL). For global fitting experiments, the working stock concentration and corresponding final assay concentration were adjusted as required. Final in-well concentrations were calculated based on measured stock concentrations and dilution factors (protein: 85/90; doxorubicin: 5/90), corresponding to an approximate final doxorubicin concentration of ~0.10 µM. Following doxorubicin addition, plates were shaken for 30 s, incubated for 2 min at room temperature, and fluorescence was measured on a BioTek Synergy Neo2 multimode plate reader using top-read optics (excitation 480/20 nm; emission 590/20 nm; gain 100; read height 8.0 mm).

Fluorescence quenching was quantified as ΔRFU, calculated as the fluorescence intensity of the corresponding doxorubicin only (no protein) control well minus the fluorescence intensity of the sample well, such that increased quenching yielded larger positive ΔRFU values. Mean ΔRFU values ± SEM were used for graphical presentation. Nonlinear regression analyses were performed using individual replicate measurements. Binding data were analyzed in GraphPad Prism 10 (GraphPad Software) with a quadratic one-site binding model accounting for ligand depletion using the following equation:

$$Y = Y_0 + (Y_{max} - Y_0) \times \left(\frac{-b - \sqrt{b^2 - (4 \times a \times c)}}{2 \times a \times L}\right)$$

Where $Y$ is ΔRFU, $Y_0$ is $\Delta RFU_{min}$, $Y_{max}$ is $\Delta RFU_{max}$, $X$ is protein concentration, $L$ is ligand concentration, $n$ is binding stoichiometry parameter, $a = 1$, $b = -K_D - \frac{L}{n} - X$, $c = \frac{L \times X}{n}$. For individual binding curves, the stoichiometry parameter was fixed at n = 1. For global analyses, datasets collected at different ligand concentrations were fit simultaneously with a shared $K_D$, while all other parameters were allowed to vary independently between datasets.

*Circular dichroism*

Circular dichroism spectra were collected on a Jasco J-810 spectropolarimeter using a 0.1 cm path length quartz cuvette. Protein samples were prepared in 1× PBS (pH 7.4) at concentrations of 10–15 µM, and concentrations were verified by UV–visible spectroscopy prior to measurement. Holo samples were prepared by addition of a two-fold molar excess of doxorubicin and left to incubate at room temperature for 30 min.

Spectra were collected from 190–250 nm in continuous scanning mode at 50 nm/min with a bandwidth of 1 nm and five accumulations per sample. Thermal denaturation experiments were performed by monitoring ellipticity at 222 nm from 20–95 °C at 2 °C temperature intervals with a heating rate of 2 °C/min.

*Monomeric fraction analysis.*

Monomeric fractions were quantified directly from the preparative size-exclusion chromatography chromatograms obtained during protein purification. Protein samples from 500 mL expression cultures were purified by preparative size-exclusion chromatography as described above, and chromatograms were exported and analyzed using GraphPad Prism 10. Baseline correction was performed by subtracting the average absorbance between 6 and 7 mL from each trace. To ensure consistent analysis across constructs, a fixed integration range of 7–18 mL was used to define the total eluted protein, and a predefined integration range of 13–16 mL was used to define the monomeric fraction. Areas under the curve (AUCs) for each region were calculated using the AUC analysis in GraphPad Prism. The percentage of monomeric protein was calculated as:

$$Monomeric\ fraction\ (\%) = 100 \times \frac{AUC_{13-16mL}}{AUC_{7-18mL}}$$

*Cell viability assay*

COV362 cells were cultured in DMEM media (Gibco, Cat# 10569010), supplemented with 10% fetal bovine serum (Corning, 35-010-CV) and 1% Pen Strep (Gibco, 15140-122). Cells were seeded in a 96-well plate at a density of 2,000 cells per well. After overnight, doxorubicin of serial 4-fold dilutions, ranging from 0 to 1,000 nmol/L, was preincubated with 20 μM doxorubicin binder proteins in media at room temperature for 5 min, and added to cells. After 7 days, cell confluence was measured using an IncuCyte Live Cell Imager (Sartorius; mean ± SD, n = 2). The values were normalized to those of cells treated by DMSO vehicle. Half maximal inhibitory concentration $(IC)_{50}$ concentrations were calculated by Nonlinear regression (curve fit) (log(inhibitor) vs. response-variable slope (four parameters)) using GraphPad Prism 10 (RRID:SCR_002798).

*X-ray crystallography*

A 3-fold molar excess of doxorubicin hydrochloride (stock solution in DMSO) was added to DoxSand2 in 1× PBS (pH 7.4) and incubated for 30 min at room temperature. Because residual free doxorubicin interfered with accurate spectrophotometric determination of the holo-protein concentration, the protein concentration was estimated from the initial apo-DoxSand2 concentration, which was determined by absorbance at 280 nm using an extinction coefficient of 13,980 $M^{-1}$ $cm^{-1}$, assuming complete protein recovery throughout the loading and concentration steps.

Following incubation, the sample was centrifuged at 21,100 × g for 10 min at room temperature to pellet any precipitated material. The clarified supernatant was then transferred to an Amicon Ultra 0.5 mL centrifugal filter (10 kDa molecular weight cutoff). The sample was concentrated by centrifugation and washed three times with 300–400 µL of 1× PBS by repeatedly concentrating the retentate and replenishing the filter with fresh buffer. This reduced the concentration of unbound doxorubicin, although residual free ligand was expected to remain. The sample was subsequently concentrated to a final estimated protein concentration of 12.5 mg/mL Crystallization conditions were screened against the Hampton JCSG+ Screen, Peg/Ion Screen, and Classics 1&2 Screens using a Mosquito Crystal instrument.

Based on these screening conditions, additional hanging drop optimization conditions were explored. The reported structure was obtained by diffraction of a ruby, hexagonally prismatic crystal grown in a hanging drop containing 500 mM HEPES, 489 mM Sodium Citrate, 68.5 mM Sodium Chloride, 5 mM Sodium phosphate dibasic, 0.9 mM Potassium phosphate monobasic, and 1.35 mM Potassium chloride at pH 7.25. Crystals were harvested and flash-frozen in liquid nitrogen without cryo protection.

*X-ray crystallography data collection and processing*

Diffraction data was collected at ALS Beamline 8.3.1 on a PILATUS3 S 6M detector with a 1.11582 Å beam wavelength at 100 K. Data were processed with XDS,[8] and statistics for data processing and structural refinement can be found in Table S3. Using the Phenix[9] suite, the structure was solved by molecular replacement with Phaser[10] using a DoxSand2 Chai-1 design model. Phenix and Servalcat[11] were used for refinement, and Elbow[12] was used for ligand restraint construction based on optimized coordinates. Cofactor placement was based on residual electron density in the binding site. When ligand occupancy was allowed to vary, full occupancy was found to best explain the data. Protein model building and adjustments were performed in Coot1[13], and MolProbity[14] was used in validation. The protein structure was deposited to the protein database as entry 36iq. All raw data were deposited to the Integrated Resource for Reproducibility in Macromolecular Crystallography[15] (proteindiffraction.org), from which they can be freely downloaded (doi:10.18430/M336IQ).

### SUPPLEMENTARY FIGURES

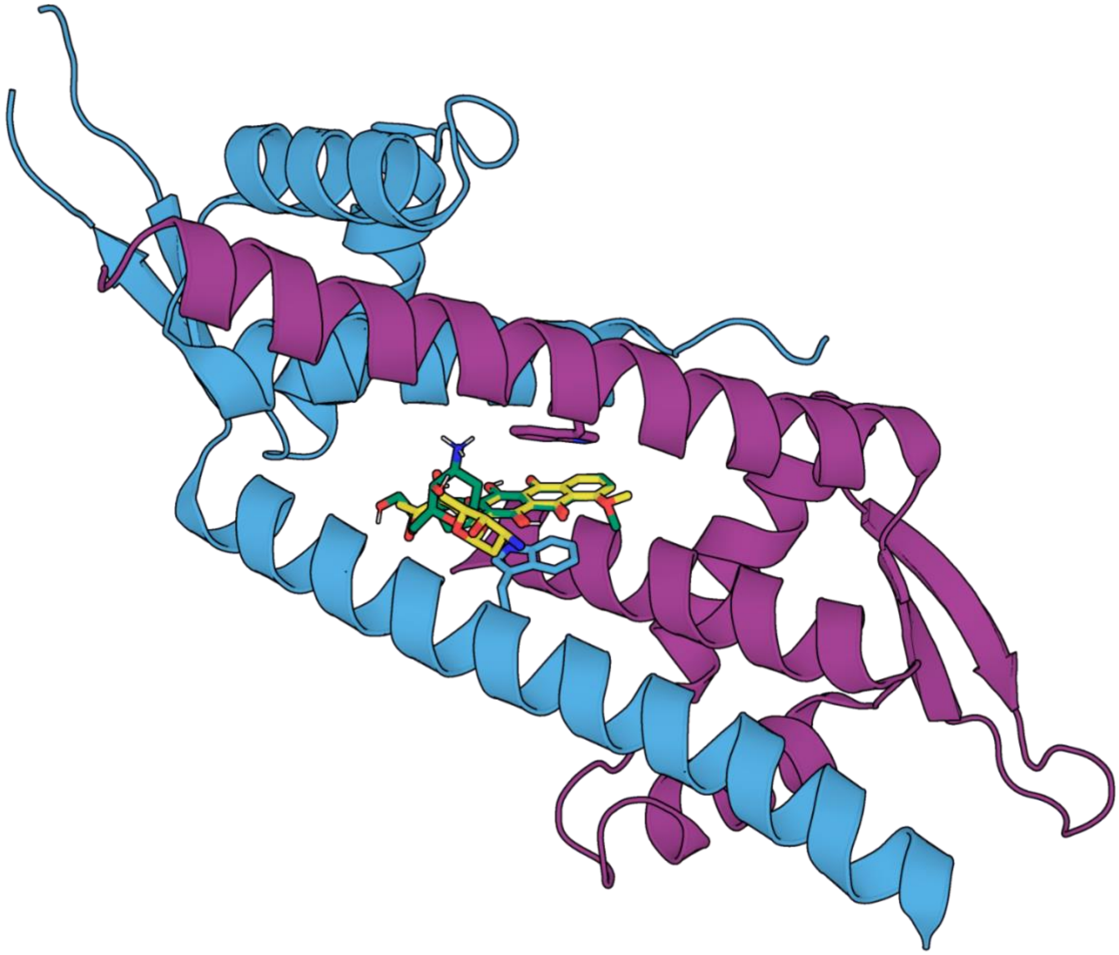

**Figure S1.** Crystal structure of the daunomycin-bound LmrR dimer (PDB: 3F8F). The protein is colored by chain (blue and magenta). Daunomycin from the crystal structure is shown as green sticks. The relative ligand pose of doxorubicin (yellow) for motif-guided protein design was defined by overlaying its anthracycline core onto the crystallographic daunomycin core.

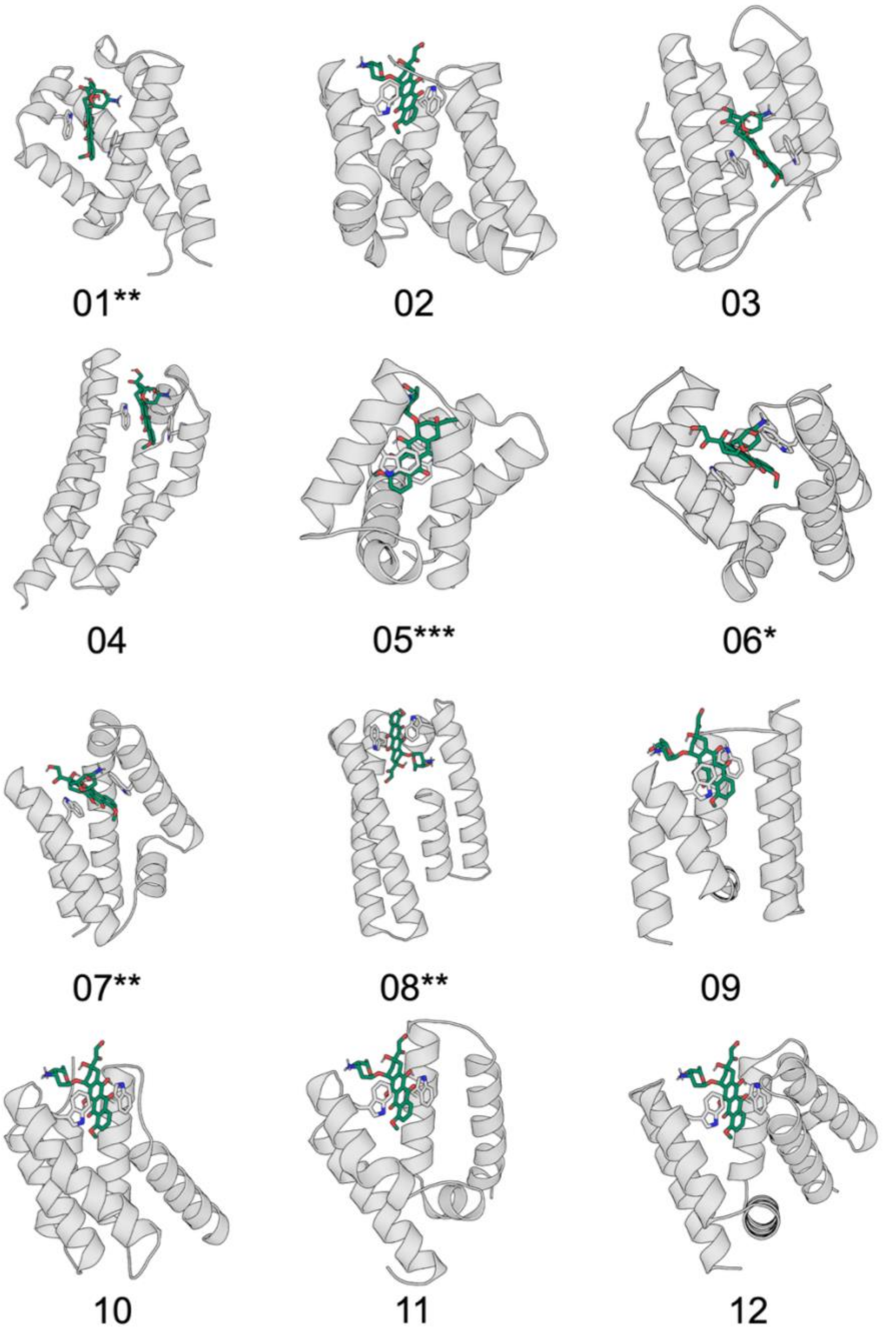


**Figure S2.** RFdiffusion All-Atom scaffold architectures selected for sequence design and experimental evaluation. Twelve backbone scaffolds generated by RFdiffusion All-Atom and advanced through the sequence design and structure-prediction pipeline are shown. Asterisks denote experimental outcomes: (*) scaffold selected for gene synthesis; (**) scaffold expressed as soluble protein; (***) scaffold expressed as soluble protein and exhibited measurable doxorubicin binding.

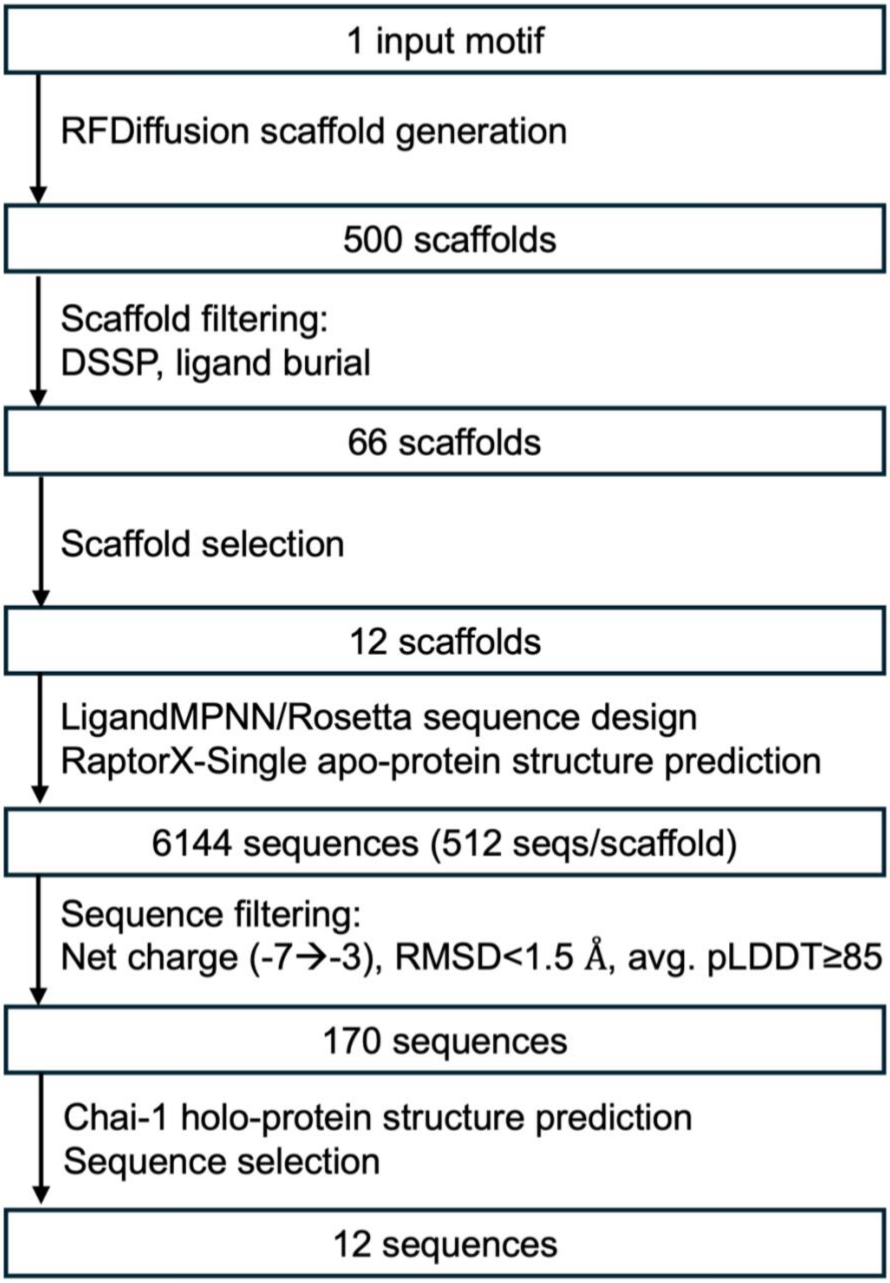


**Figure S3.** Progressive filtering of the computational design search space. A six-residue aromatic π-stacking motif derived from the daunomycin-bound LmrR crystal structure (PDB: 3F8F) was provided as the fixed input motif to RFdiffusion All-Atom. Sequential filtering based on scaffold quality, ligand burial, secondary structure content, sequence design, and structure prediction progressively reduced the design space to 12 sequences representing six scaffold architectures that were selected for experimental characterization. Each box indicates the design entity and the number carried forward to the next stage.

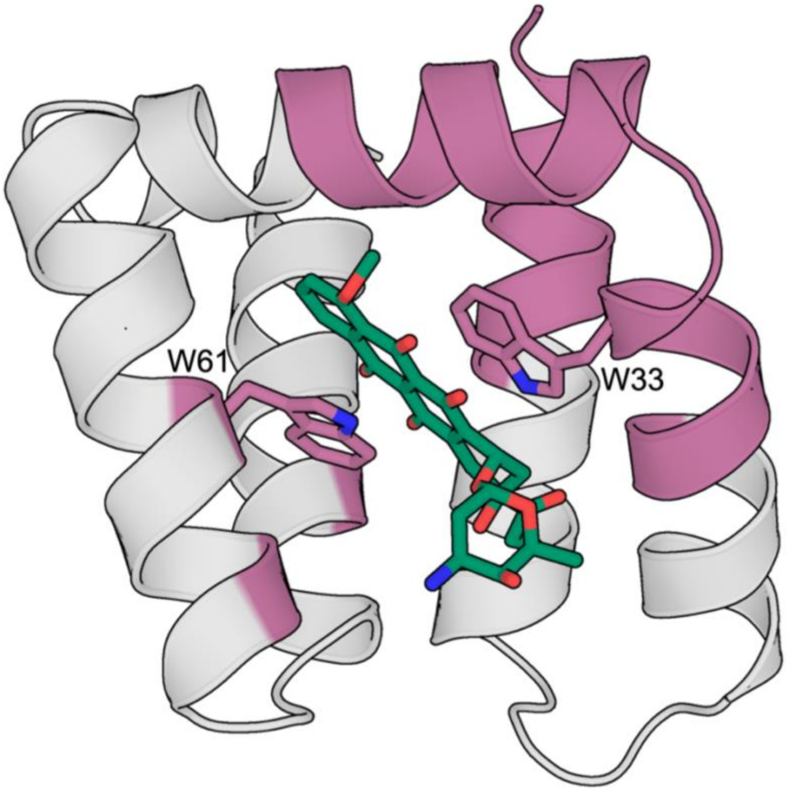


**Figure S4.** DoxSand0 scaffold after loop optimization using MASTER. Residues held fixed during sequence design (31% of the scaffold) are highlighted in pink, while the remaining residues were allowed to vary. Doxorubicin is shown as green sticks. Residues W33 and W61, shown as sticks, were constrained relative to doxorubicin using Rosetta AtomPair constraints during sequence design to preserve the binding-site geometry.

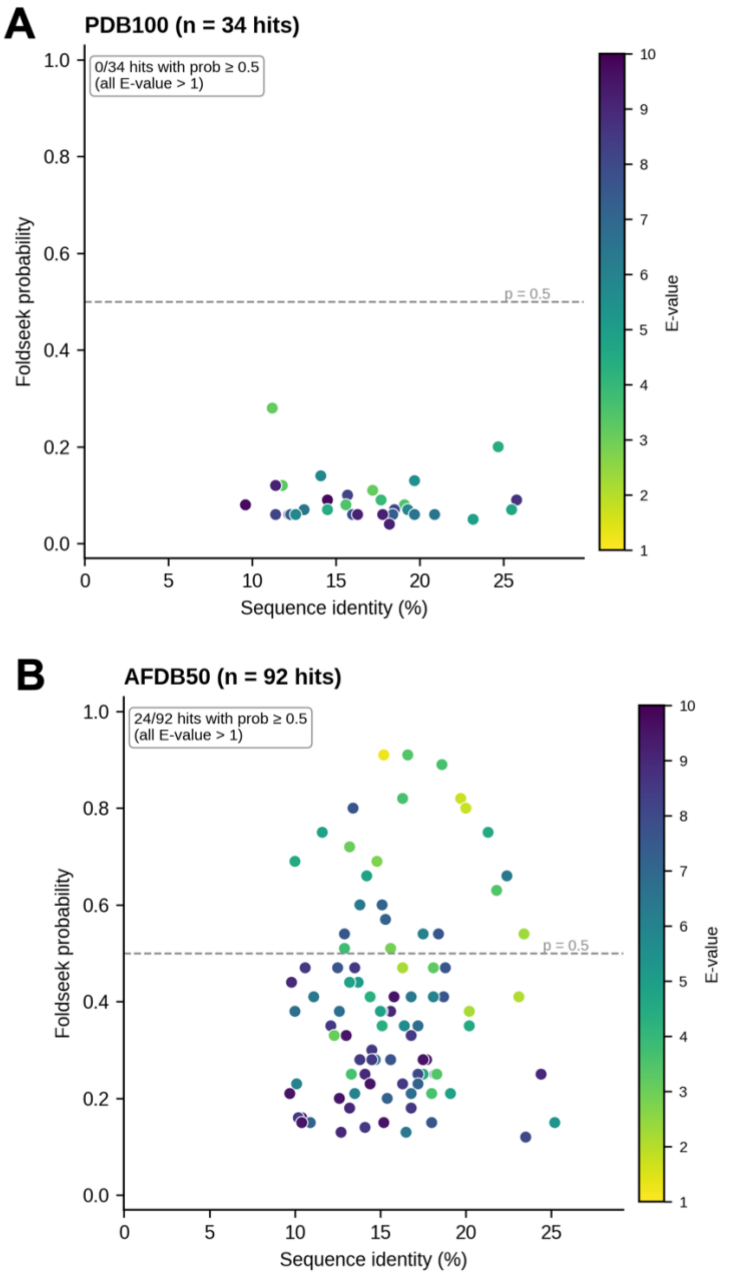


**Figure S5.** Foldseek (3Di/AA mode) searches against the PDB100 (A) and AlphaFold/UniProt50 (B) databases identified only weak structural matches. Sequence identity is plotted against Foldseek probability, with point color indicating E-value.[16,17] No statistically significant structural homologs were identified in either database (all E-values > 1).

### SUPPLMENTARY TABLES

**Table S1.** De novo Small-Molecule Binding Proteins with Experimentally Determined Holo Structures

| protein | ligand | sequence length | scaffold | $K_D$ (nM) |
|---|---|---|---|---|
| DoxSand2 | doxorubicin | 85 | mini helical jaw | 85 |
| mFAP1[18] | DFHBI | 112 | β-barrel | 560 |
| A1E[19] | progesterone | 113 | NTF2 fold | 5000-10000 |
| apx1049[20] | apixaban | 119 | NTF2 fold | 650 |
| ABLE[21] | apixaban | 126 | 4-helix bundle | 5000 |
| hcy129_mpnn5[20] | cortisol | 135 | NTF2 fold | 2300 |
| PiB[22] | rucaparib | 147 | 4-helix bundle | <5 |
| EPIC Q51N[23] | exatecan | 148 | 4-helix bundle | 8 |

**Table S2.** Protein and DNA Sequences of DoxSand Variants

| Protein | Protein sequence | DNA sequence |
|---|---|---|
| DoxSand0 | MEKEIVEEALAILK<br>AVLDGEPGAEER<br>AAAFWANPENRK<br>VVTDVVVKYANKL<br>GLTKGSTDPAQAV<br>KDWWAQADAAG<br>DVAAGNAVGVKA<br>LELELENLYFQSH<br>HHHHH* | CTTTAAGAAGGAGATATACCATGGATGGAGAAGG<br>AAATAGTGGAAGAAGCTCTGGCGATCCTGAAGGC<br>GGTGCTTGATGGCGAACCTGGAGCAGAGGAGCG<br>TGCCGCAGCCTTCTGGGCAAACCCCGAAAACCG<br>AAAGGTCGTTACCGACGTAGTGGTTAAGTACGCG<br>AACAAACTTGGCTTAACTAAGGGCTCTACGGACC<br>CAGCCCAAGCCGTTAAGGACTGGTGGGCCCAGG<br>CAGATGCCGCCGGTGACGTGGCCGCGGGTAATG<br>CGGTAGGTGTAAAAGCTTTGGAGTTAGAGCTTGA<br>GAACCTGTACTTTCAAAGTCATCACCACCATCAC<br>CACTGACTCGAGCACCACCACCACCACCAC |
| DoxSand1 | MEKEIVEEALKLV<br>QGFLDDPNDKAV<br>LEAAAAFWANPE<br>NRKVVTDTVAKEL<br>GISSEELEARWRE<br>YDAAGRLAEHNEI<br>VAKGLRKALENLY<br>FQSHHHHHH* | CTTTAAGAAGGAGATATACCATGGATGGAAAAGG<br>AGATTGTGGAAGAGGCTCTGAAGCTGGTGCAGG<br>GCTTTCTCGACGACCCGAACGACAAGGCAGTGC<br>TCGAAGCCGCGGCCGCGTTCTGGGCCAATCCAG<br>AGAACCGTAAAGTGGTTACCGATACAGTAGCGAA<br>GGAGTTAGGTATTTCGTCCGAGGAGCTTGAGGC<br>CCGCTGGCGCGAGTACGACGCAGCAGGCAGATT<br>AGCAGAACATAATGAGATCGTCGCGAAGGGTTTA<br>CGTAAGGCCCTCGAAAATCTGTACTTCCAATCTC<br>ATCATCACCACCATCATTAGCTCGAGCACCACCA<br>CCACCACCAC |
| DoxSand2 | MEKEIVEEALKLV<br>QGFLDDPNDKAV<br>LEAAAAFWANPE<br>NRKVVTDTIAKEL<br>GISSEELEARWRE<br>YDAAGRLAEANEI<br>VAKGLRKALENLY<br>FQSHHHHHH* | CTTTAAGAAGGAGATATACCATGGATGGAGAAAG<br>AAATCGTTGAAGAGGCCCTTAAGTTAGTCCAAGG<br>CTTCCTGGACGATCCAAACGATAAGGCAGTTCTG<br>GAAGCTGCAGCTGCATTCTGGGCAAACCCTGAA<br>AATCGGAAGGTAGTAACGGACACGATCGCCAAAG<br>AACTGGGTATCTCAAGCGAGGAATTAGAGGCGC<br>GCTGGAGAGAGTATGATGCCGCTGGCCGACTCG<br>CCGAGGCAAACGAGATCGTCGCGAAGGGCCTTC<br>GCAAGGCACTGGAAAATTTGTATTTCCAGTCACAT<br>CACCATCACCATCACTGACTCGAGCACCACCACC<br>ACCACCAC |

**Table S3.** Crystallographic data processing and refinement statistics for holo DoxSand2.

| | **Value** |
|---|---|
| PDB ID | 36IQ |
| Space group | P 32 2 1 |
| Cell constants<br>a, b, c, α, β, γ | 42.70 Å, 42.70 Å, 174.71 Å<br>90.00°, 90.00°, 120.00° |
| Resolution (Å) | 58.24 – 1.56 |
| % Data completeness | 99.7 (58.24-1.56) |
| Rmerge | 0.18 |
| < *I/σ(I)* > | 1.56 |
| R, $R_{free}$ | 0.218, 0.255 |
| $R_{free}$ test set | 1999 reflections (6.77%) |
| Wilson B-factor (Å$^2$) | 25.1 |
| Anisotropy | 0.256 |
| Bulk solvent<br>$k_{sol}$(e/Å$^3$), $B_{sol}$(Å$^2$) | 0.37, 28.6 |
| L-test for twinning | < \|L\| > = 0.47, < L2 > = 0.30 |
| Estimated twinning fraction | 0.046 for -h,-k,l |
| Fo,Fc correlation | 0.95 |
| Total number of atoms | 3071 |
| Average B, all atoms (Å$^2$) | 38.0 |
| Protein residues | 180 |
| RMS (bonds) | 0.007 |
| RMS (angles) | 0.79 |
| Ramachandran favored (%) | 97.73 |
| Ramachandran allowed (%) | 2.27 |
| Ramachandran outliers (%) | 0.00 |
| Rotamer outliers (%) | 0.69 |
| Clashscore | 2.33 |

### SUPPLEMENTARY TEXTS

#### Quantum chemical calculations

Density functional theory optimization of the cationic doxorubicin geometry was carried out in ORCA 6.1.1[24,25] using the B3LYP functional with a D3BJ[26,27] dispersion correction and def2-SVP set. The reported structure was concluded to be a ground-state minimum based on the analysis of the harmonic vibrational analytical frequencies, which show no imaginary frequencies.

Doxorubicin coordinates:

| | | | |
|---|---|---|---|
| C | -21.09693843452550 | -2.30349715515109 | 10.72262167574020 |
| C | -20.79468975060255 | -3.47656711563832 | 9.81163732826365 |
| O | -21.93740181405846 | -3.67670252351532 | 8.96019234677085 |
| C | -20.52628542018282 | -4.78202881835712 | 10.56580332752132 |
| O | -21.60057538733238 | -5.14578753810384 | 11.41999901088420 |
| C | -20.28503823548590 | -5.91126567344913 | 9.54859530443355 |
| N | -20.15898426480298 | -7.18846789587201 | 10.32245821844999 |
| C | -21.44339779655918 | -6.01880372649053 | 8.56157005024540 |
| C | -21.72602347213343 | -4.64914158043469 | 7.94500059353509 |
| C | -17.13629240435094 | -6.71522001278077 | -0.11058125414148 |
| C | -17.41810963911604 | -7.89307292240745 | -0.80041166549987 |
| C | -18.49779491646132 | -8.68900301891866 | -0.42914869070723 |
| C | -19.32787751343310 | -8.32296837041791 | 0.64988362955414 |
| O | -20.36937901965412 | -9.07049656740681 | 1.02937907414792 |
| C | -19.05772593494920 | -7.11927040524125 | 1.36758241153209 |
| C | -19.89627058438804 | -6.67569154398671 | 2.50374477736919 |
| O | -20.88415538240861 | -7.32586383517695 | 2.89678060116937 |
| C | -19.54350432689929 | -5.42170851453473 | 3.18385140155990 |
| C | -20.33161887466574 | -4.98129221758535 | 4.26396619832619 |
| O | -21.39559090172265 | -5.66468640513066 | 4.68274230877933 |
| C | -20.00974279013777 | -3.76158753785334 | 4.93182634296934 |
| C | -20.88045718860018 | -3.36128770012380 | 6.09933419591702 |
| O | -20.64718718051054 | -4.32996081991562 | 7.12598242060150 |
| C | -20.65042918723356 | -1.92109265769042 | 6.55586272295201 |
| C | -19.16477730860662 | -1.53297675611269 | 6.59394261030909 |
| O | -18.52010506560866 | -2.36868795370375 | 7.54631242507089 |
| C | -19.09218527590996 | -0.06596615609807 | 7.03283908419579 |
| O | -19.39543509536898 | 0.84766248717994 | 6.28973571582890 |
| C | -18.67878163364835 | 0.23691192791372 | 8.45729157198461 |
| C | -18.55708748445511 | -1.69812967415249 | 5.19856936578319 |
| C | -18.92166146893204 | -2.99847968292543 | 4.53395214438609 |

| | | | |
|---|---|---|---|
| C | -18.12149667323065 | -3.43549841253278 | 3.43706579724489 |
| C | -18.42968617580821 | -4.63709756210853 | 2.77310215605001 |
| C | -17.60043948984298 | -5.06497105977099 | 1.65630573888087 |
| O | -16.62144344509488 | -4.39051364490054 | 1.28414729078038 |
| C | -17.94653717286498 | -6.33133904456175 | 0.96106942040707 |
| C | -20.67372187513569 | -10.27435532789593 | 0.34049131345024 |
| H | -19.28356150608560 | -7.22508471806262 | 10.86373559502561 |
| H | -20.94198089165320 | -7.23834899491197 | 10.99580722250825 |
| H | -17.56235260578633 | -2.30970594895921 | 7.40774604471369 |
| H | -22.42467038692296 | -4.86905827631136 | 10.98509377408131 |
| H | -21.44333331461864 | -6.46626988899691 | 4.07252465369209 |
| H | -16.29526678625109 | -6.08007061869547 | -0.38844157712256 |
| H | -16.79075337651188 | -8.20026610795967 | -1.64063322894662 |
| H | -18.69881827187165 | -9.60468395199494 | -0.98296597788819 |
| H | -22.66634457667861 | -4.66990278039658 | 7.36944952760049 |
| H | -21.93539372107437 | -3.45173687441778 | 5.78651877257764 |
| O | -17.08937757923668 | -2.67547956430412 | 3.07398945985512 |
| H | -19.33668358451593 | -5.74905817441032 | 9.02010600957075 |
| H | -19.63314977973288 | -4.66260837459468 | 11.20056701617475 |
| H | -19.91454407679745 | -3.25077815583157 | 9.18930411087640 |
| H | -21.11889253035856 | -1.77416226814696 | 7.53786236148043 |
| H | -21.21754412497226 | -6.73498436259120 | 7.75834300944037 |
| H | -21.15237170813612 | -1.24413582418294 | 5.84829657597909 |
| H | -22.35255159504149 | -6.36138794503422 | 9.08061326066983 |
| H | -19.30975036115986 | -0.37141992389533 | 9.13430189110308 |
| H | -17.64416369701893 | -0.14013995576244 | 8.59570197216914 |
| O | -18.77975248220344 | 1.59780582206059 | 8.73254764448641 |
| H | -18.89280378177246 | -0.85694430015677 | 4.56931338282568 |
| H | -17.46025756198830 | -1.60353240812577 | 5.25340850524468 |
| H | -20.91488060876335 | -10.08637422353297 | -0.71944178347139 |
| H | -19.84512803566693 | -10.99991456095369 | 0.40425359911444 |
| H | -21.55700840312724 | -10.69358690685340 | 0.84010025561144 |
| H | -21.30298024662072 | -1.40439225382596 | 10.12333578537467 |
| H | -20.23344192827501 | -2.09626420140046 | 11.37316656101464 |
| H | -21.97117186679913 | -2.50331327673373 | 11.36065798440931 |
| H | -16.65881632464369 | -3.14977755969153 | 2.30262384451004 |
| H | -20.19149697322086 | -8.02287999087298 | 9.72048227713987 |
| H | -19.04538316747202 | 2.01367622569897 | 7.89130600673360 |

**Doxorubicin ligand Rosetta parameters files (DM2_params_file):**

```
NAME DM2
IO_STRING DM2 Z
TYPE LIGAND
AA UNK
ATOM  C15 aroC  X   -0.02
ATOM  C14 aroC  X   -0.02
ATOM  O5  OH    X   -0.56
ATOM  H12 Hpol  X   0.53
ATOM  C13 aroC  X   -0.02
ATOM  C12 aroC  X   -0.02
ATOM  O4  OOC   X   -0.66
ATOM  C11 aroC  X   -0.02
ATOM  C10 aroC  X   -0.02
ATOM  O3  OH    X   -0.56
ATOM  C27 CH3   X   -0.17
ATOM  H27 Hapo  X   0.19
ATOM  H28 Hapo  X   0.19
ATOM  H29 Hapo  X   0.19
ATOM  C9  aroC  X   -0.02
ATOM  C8  aroC  X   -0.02
ATOM  C7  aroC  X   -0.02
ATOM  C26 aroC  X   -0.02
ATOM  C25 aroC  X   -0.02
ATOM  O11 OOC   X   -0.66
ATOM  C24 aroC  X   -0.02
ATOM  C23 aroC  X   -0.02
ATOM  O10 OH    X   -0.56
ATOM  H19 Hpol  X   0.53
ATOM  C22 aroC  X   -0.02
ATOM  C21 CH2   X   -0.08
ATOM  C18 CH1   X   0.01
ATOM  O7  OH    X   -0.56
ATOM  H15 Hpol  X   0.53
ATOM  C17 CH2   X   -0.08
ATOM  C16 CH1   X   0.01
ATOM  O6  OH    X   -0.56
ATOM  C6  CH1   X   0.01
ATOM  O1  OH    X   -0.56
ATOM  C2  CH1   X   0.01
ATOM  C1  CH3   X   -0.17
ATOM  H1  Hapo  X   0.19
ATOM  H2  Hapo  X   0.19
ATOM  H3  Hapo  X   0.19
```

**Rosetta constraints (constraints_file) used for re-design of DoxSand0**

```
AtomPair NE1 60A C12 1X HARMONIC 3.4 0.2
AtomPair CH2 60A C23 1X HARMONIC 3.1 0.2
AtomPair CG 60A C10 1X HARMONIC 4.1 0.2
AtomPair CD2 60A C26 1X HARMONIC 4.0 0.2
AtomPair CZ2 60A C22 1X HARMONIC 3.5 0.2
AtomPair CD2 32A C14 1X HARMONIC 3.8 0.2
AtomPair CZ2 32A C24 1X HARMONIC 3.2 0.2
AtomPair CD1 32A C15 1X HARMONIC 3.8 0.2
AtomPair CH2 32A C25 1X HARMONIC 3.5 0.2
```
